# Revealing the Organ-Specific Expression of *SPTBN1* using Single-Cell RNA Sequencing Analysis

**DOI:** 10.1101/2023.06.01.543198

**Authors:** Jongyun Jung, Qing Wu

## Abstract

Despite the recent technological advances in single-cell RNA sequencing, it is still unknown how three marker genes (*SPTBN1*, *EPDR1*, and *PKDCC*), which are associated with bone fractures and highly expressed in the muscle tissue, are contributing to the development of other tissues and organs at the cellular level. This study aims to analyze three marker genes at the single-cell level using 15 organ tissue types of adult human cell atlas (AHCA). The single-cell RNA sequencing analysis used three marker genes and a publicly available AHCA data set. AHCA data set contains more than 84,000 cells from 15 organ tissue types. Quality control filtering, dimensionality reduction, clustering for cells, and data visualization were performed using the Seurat package. A total of 15 organ types are included in the downloaded data sets: Bladder, Blood, Common Bile Duct, Esophagus, Heart, Liver, Lymph Node, Marrow, Muscle, Rectum, Skin, Small Intestine, Spleen, Stomach, and Trachea. In total, 84,363 cells and 228,508 genes were included in the integrated analysis. A marker gene of *SPTBN1* is highly expressed across all 15 organ types, particularly in the Fibroblasts, Smooth muscle cells, and Tissue stem cells of the Bladder, Esophagus, Heart, Muscle, Rectum, Skin, and Trachea. In contrast, *EPDR1* is highly expressed in the Muscle, Heart, and Trachea, and *PKDCC* is only expressed in Heart. In conclusion, *SPTBN1* is an essential protein gene in physiological development and plays a critical role in the high expression of fibroblasts in multiple organ types. Targeting *SPTBN1* may prove beneficial for fracture healing and drug discovery.

**Author Summary:** Three marker genes (*SPTBN1*, *EPDR1*, and *PKDCC*) are playing a critical role in the shared genetic mechanisms between bone and muscle. However, how these marker genes contribute to developing other tissues and organs at the cellular level is unknown. Here, we build on prior work to study an unappreciated degree of heterogeneity of three marker genes in 15 adult human organs by using single-cell RNA sequencing technology. Our analysis included 15 organ types: Bladder, Blood, Common Bile Duct, Esophagus, Heart, Liver, Lymph Node, Marrow, Muscle, Rectum, Skin, Small Intestine, Spleen, Stomach, and Trachea. In total, 84,363 cells from 15 different organ types were included. In all 15 organ types, *SPTBN1* is highly expressed, including fibroblasts, smooth muscle cells, and skin stem cells of the bladder, esophagus, heart, muscles, and rectum. The first-time discovery of the high expression of *SPTBN1* in 15 organ types suggests that it may play a critical role in physiological development. Our study concludes that targeting *SPTBN1* may benefit fracture healing and drug discovery.

## Introduction

Bone fractures represent a global public health concern, contributing to work absences, disability, reduced quality of life, health complications, and increased healthcare costs, affecting individuals, families, and societies [1]. Genetics plays a vital role in bone fractures [2], and numerous genetic variants have been associated with bone fractures in genome-wide association studies (GWAS) [3]. The recent advance of new sequencing technologies has found an important link between bone and muscle [4], suggesting that understanding these tissues’ relationships could provide insights into preventing and treating bone fractures.

Single-cell RNA sequencing (scRNA-seq) allows the characterization of transcriptomes at the single-cell level at various development stages, assessing distinct cell types and states, their dynamic trajectories, and molecular programs governing sequential cell fates in skeletal muscle development [5]. A recent study by He et al. [6] examined the single-cell transcriptomes of 84,363 cells derived from 15 organ types of one adult donor and generated an adult human cell atlas. They found the inter- and intra-organ heterogeneity of cell characteristics during the development of human diseases.

Our previous study [7] found that three marker genes (*SPTBN1*, *EPDR1*, and *PKDCC*) from GWAS findings play a critical role in muscle tissue, which has implications for bone fracture risk. However, how these marker genes contribute to developing other tissues and organs at the cellular level is unknown. To fully understand the developmental process of skeletal muscle and its relationship to bone health, it is essential to understand each cell type’s functional capacities and response. Despite the technological advances in scRNA-seq, limited knowledge exists in understanding the cellular-level relationship between bone and muscle. Based on previous GWAS findings involving bone fracture-related variants of three marker genes and the adult human cell atlas generated by He et al. [6], our study aims to analyze these three marker genes (*SPTBN1*, *EPDR1*, and *PKDCC*) at the single-cell level in 15 adult human organs.

## Results

### Single-cell RNA sequencing of 15 organ samples

We analyzed a total of 15 organ types in the downloaded datasets: Bladder, Blood, Common Bile Duct, Esophagus, Heart, Liver, Lymph Node, Marrow, Muscle, Rectum, Skin, Small Intestine, Spleen, Stomach, and Trachea. The raw dataset contained 251,469 genes and 84,363 cells. After quality control, the number of features was reduced to 228,508 (**Table 1**). We did not perform feature filtering on the datasets to include as many features as possible. Violin plots were used to display the results of features after quality control for each organ type (**Figure S1**).

**Table 1.** Summary of the number of genes and cell data used in this study, published by He et al. (2020) [6]. QC, Quality Control

| Organ Type | Number of genes | Number of cells | Number of genes (after QC) |
| --- | --- | --- | --- |
| Bladder | 17059 | 7572 | 16478 |
| Blood | 14552 | 1407 | 11758 |
| Common bile duct | 19042 | 3160 | 16381 |
| Esophagus | 17539 | 9117 | 17172 |
| Heart | 16473 | 7881 | 16075 |
| Liver | 17640 | 2839 | 14191 |
| Lymph Node | 14607 | 7771 | 13508 |
| Marrow | 16122 | 3230 | 13338 |
| Muscle | 16024 | 5732 | 15045 |
| Rectum | 17834 | 6280 | 16860 |
| Skin | 17773 | 7710 | 16984 |
| Small Intestine | 16412 | 4312 | 14578 |
| Spleen | 15321 | 4512 | 13496 |
| Stomach | 16566 | 5318 | 15002 |
| Trachea | 18505 | 7522 | 17642 |
| **Total** | **251469** | **84363** | **228508** |

### UMAP visualization of single-cell RNA sequencing data

Single-cell RNA sequencing data of 15 organ tissues from a male adult donor were plotted using UMAP (**Figure 2**). Each dot represents one cell, with colors coded according to the organ of origin. UMAP visualization of all cells (84,363) in 15 organs and tissues shows distinct clusters of organ types for Heart, Bladder, Blood, Esophagus, Lymph, and Liver (**Figure 2A**). A total of 27 clusters were identified using UMAP visualization (**Figure 2B**).

**Figure 1.**
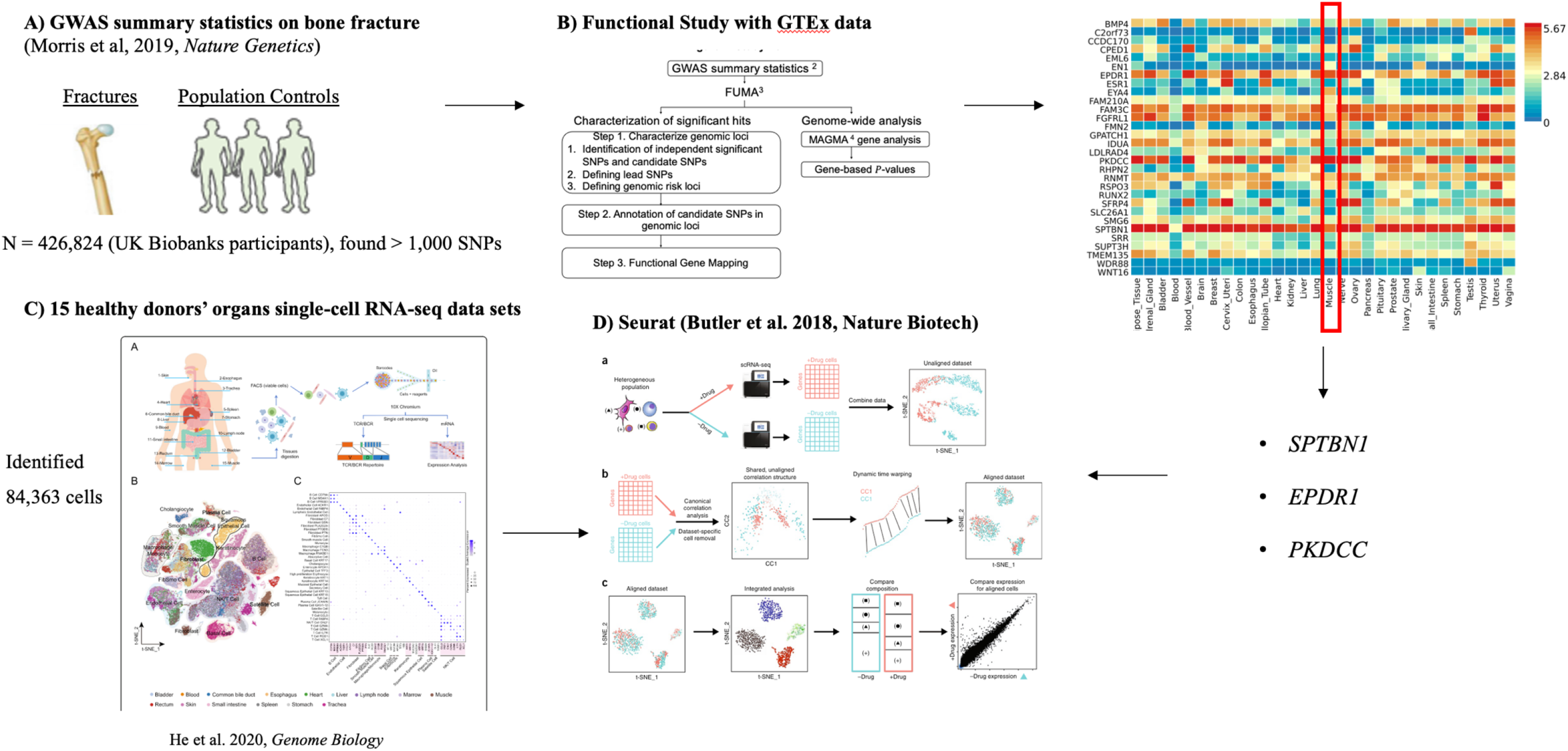
The workflow of integrative genomic analysis. **A** Genome-Wide Association Study (GWAS) summary statistics on bone fractures was used [3]. **B** Functional study based on the functional mapping and annotations of GWAS findings were utilized to prioritize the causal Single-Nucleotide Polymorphisms using the Genotype-Tissue Expression Data. **C** He et al. (2020) [6] identified 84,363 cells using 15 healthy man donor’s organs of single-cell RNA sequencing data. **D** Seurat package [18] (version 4.1.0; https://satijalab.org/Seurat) was employed to perform the single-cell RNA seq data analysis.

**Figure 2.**
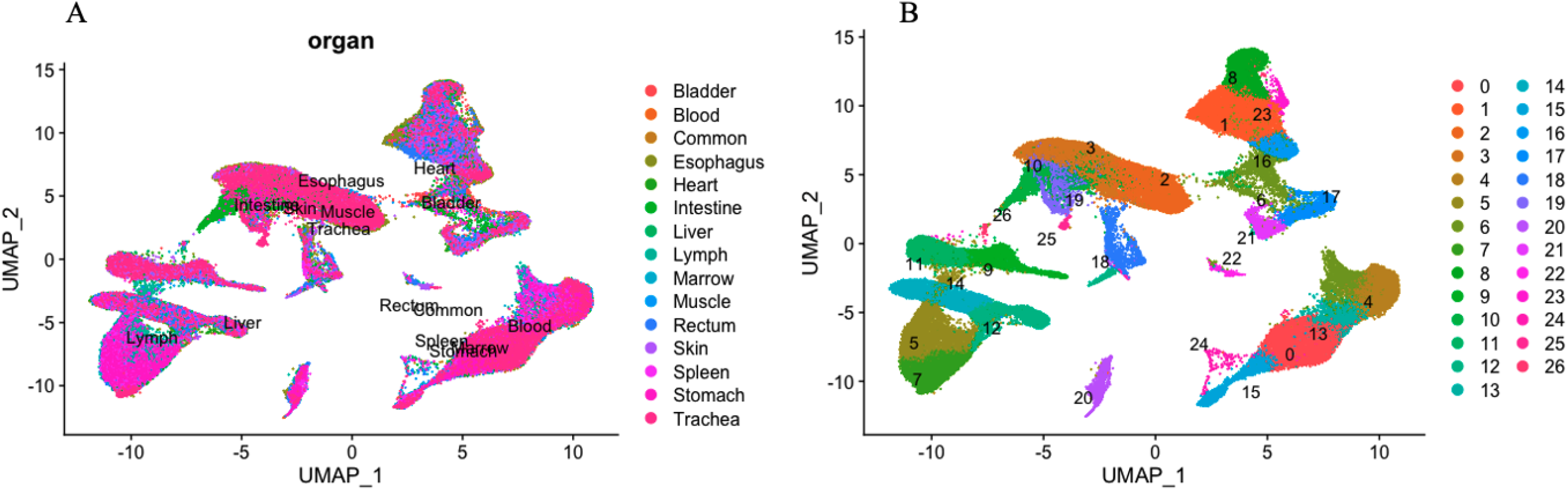
Single-cell RNA sequencing of 15 organ tissues from a male adult donor was plotted using the uniform manifold approximation and projection (UMAP). **A** UMAP visualization of all cells (84,363) in 15 organ tissues. Each dot represents one cell, with colors coded according to the origin of an organ. **B** UMAP visualization of 27 cell clusters.

### Expression of *EPDR1*, *PKDCC*, and *SPTBN1* in 15 organ types

Violin plots of the normalized expression of marker genes *EPDR1*, *PKDCC*, and *SPTBN1* are shown in **Figure 3**. Each candidate gene expression result was analyzed by 15 organ types. *SPTBN1* is highly expressed in all organ types, while *EPDR1* shows high expression in the Muscle, Heart, and Trachea. In contrast, *PKDCC* is only highly expressed in Heart. Each candidate gene expression is also analyzed by cell type in each organ (**Figure 4**, **Figure S2**, and **S3**). Despite *SPTBN1* being highly expressed in every organ, not all cell types express it. *SPTBN1* is highly expressed in the Fibroblasts, Smooth muscle cells, and Tissue stem cells of the Bladder, Esophagus, Heart, Muscle, Rectum, Skin, and Trachea. It is also highly expressed in the B-cell and T-cells of the Lymph node, Spleen, and Marrow. In contrast, *PKDCC* is only expressed in the Fibroblasts of the Bladder and Rectum. *EPDR1* is expressed in the Tissue stem cells of Heart, and Muscle. The average expression of marker genes *SPTBN1*, *EPDR1*, and *PKDCC* in each cluster was plotted using the dot plot in **Figure 5**. *SPTBN1* is highly expressed (> 60% expression) in clusters from the Heart (clusters 1 and 8), muscle (clusters 9, 11, and 26), Bladder (cluster 16), and Rectum (cluster 22). However, *PKDCC* and *EPDR1* do not express highly in any of the clusters.

**Figure 3.**
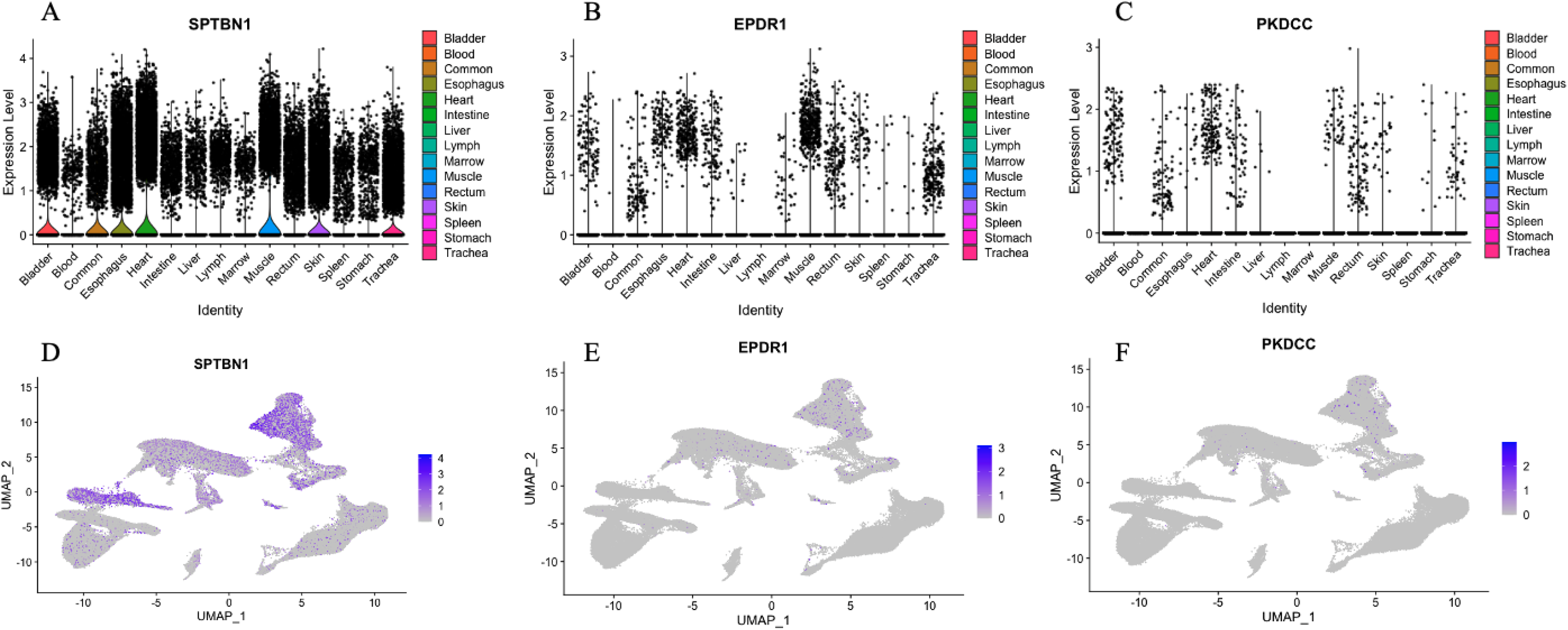
Violin plots of the normalized expression of marker genes for **A**: *SPTBN1*, **B**: *EPDR1*, and **C**: *PKDCC*. For each panel, the y-axis shows the normalized expression level for a marker gene, as indicated in the title, and the x-axis indicates the organ types. UMAP visualization of the normalized expression marker genes for **D**: *SPTBN1*, **E**: *EPDR1*, and **F**: *PKDCC*. Each dot represents one cell, with a color from gray to blue representing the expression level from low to high.

**Figure 4.**
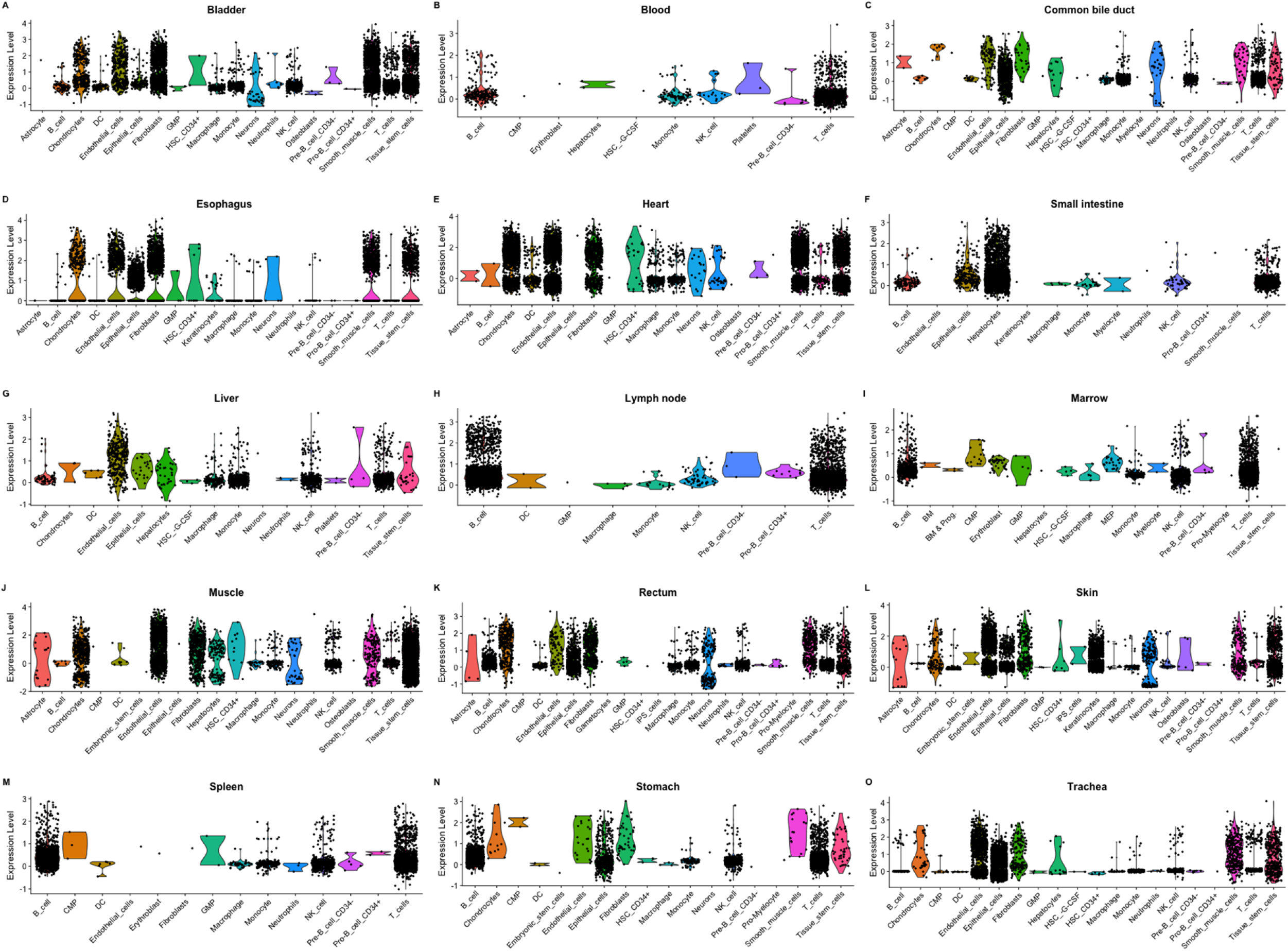
Violin plots of the normalized expression of *SPTBN1* for each organ by the cell type.

**Figure 5.**
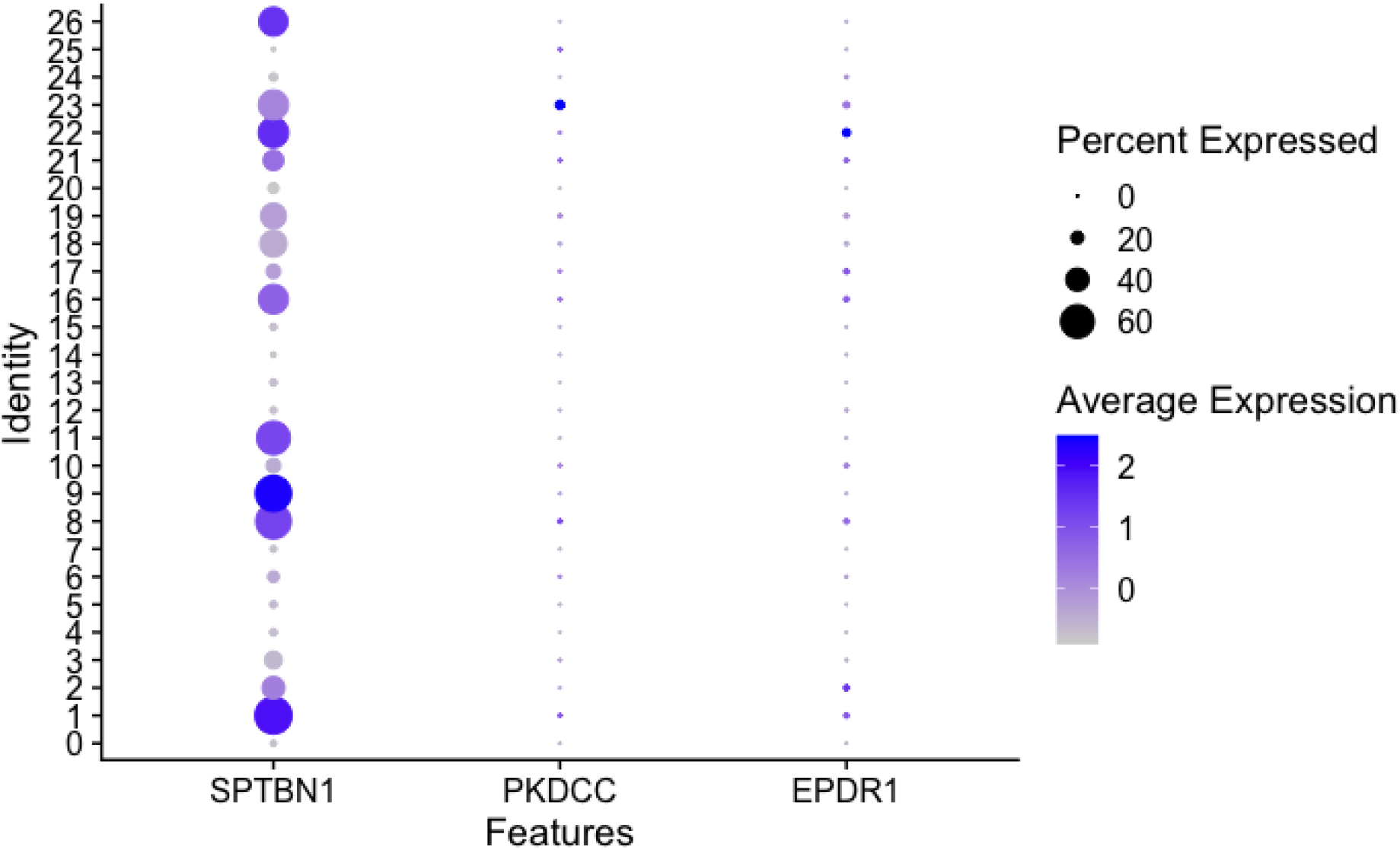
Dot plot of the average expression of marker genes for *SPTBN1 EPDR1*, and *PKDCC* in each cluster (identity). Each marker gene was grouped by each cluster. Circle size indicates the percent expressed in each cluster, and color represents the average expression level from low to high.

## Discussion

Through single-cell RNA sequencing analysis, our study offers valuable insights into the role of GWAS-identified marker genes in various organ types. We used the largest high-resolution adult human cell atlas of single-cell RNA datasets to examine the expression of key marker genes in 15 different organ types. Our findings revealed that *SPTBN1* is highly expressed in all 15 organ types, including fibroblasts, smooth muscle cells, and skin stem cells of the bladder, esophagus, Heart, muscles, and rectum. This first-time discovery of the high expression of *SPTBN1* in 15 organ types suggests that it may play a critical role in physiological development.

Our results are consistent with recent studies. For example, a study by Yang et al. (2021) [8] found that *SPTBN1* is a cytoskeleton protein in all nucleated cells, contributing to organ development by establishing and maintaining cell structure and regulating various cellular functions. They also found that *SPTBN1* involves bone structure development and fracture healing. Another study by Cousin et al. (2021) [9] identified *SPTBN1* as a key player in brain and central nervous system development, with *SPTBN1* variant carriers presenting a neurodevelopmental syndrome. Comparing wild-type and knock-out mice, they observed clear differences in physiological development. A previous study conducted by Xu et al. [10] showed that *SPTBN1* prevents primary osteoporosis development by modulating VEGF, TGF-beta/Smad3, and STAT1/Cxcl9 signaling pathways.

By using the scRNA-seq technology with data from several organ types, our study has revealed an unappreciated degree of heterogeneity in fibroblasts within and across organ types. Fibroblasts are critical in tissue development and repair mechanisms [11]. Recent multiple scRNA-seq studies show that fibroblasts display transcriptional changes similar to cellular differentiation trajectories during tissue development and repair [12,13]. Previous studies have shown the unexpected similarities and unique characteristics of fibroblasts across diverse organs such as the skin, lungs, Heart, and skeletal muscle that are currently being leveraged for treating human diseases. We find that fibroblasts exhibit high expression in the bladder, rectum, and esophagus in addition to previously known organ types. To our knowledge, none of the previous studies found a high expression of fibroblasts in these organ types. Furthermore, our study revealed that *SPTBN1* is a critical gene in the high expression of fibroblasts in multiple organ types.

Our study has several limitations. First, although we utilized a large adult human cell atlas of single-cell RNA sequencing data, comprising 15 organ types and over 84,000 cells, the dataset lacks representation of other organ types that may play a critical role in physiological development. Second, we only tested three marker genes highly expressed in muscle tissue, as identified in our previous study. Other genes not highly expressed in muscle tissue may also play important roles. Lastly, we have not conducted trajectory analysis to investigate the origin of SPTBN1 genes in these datasets.

In conclusion, our study demonstrates that *SPTBN1* is highly expressed in all organ types, *EPDR1* in muscles, Heart, and trachea, and *PKDCC* in the Heart. These findings suggest that *SPTBN1* is an essential protein gene in physiological development. Targeting *SPTBN1* may prove beneficial for fracture healing and drug discovery.

## Materials and Methods

### Genome-Wide Association Study (GWAS) Summary Dataset

Morris et al. [3] conducted a comprehensive GWAS investigating genetic variants of bone mineral density (BMD) as estimated by heel quantitative ultrasound (eBMD) in 426,824 UK Biobank participants. They identified 518 genome-wide BMD-significant loci and 13 bone fracture loci (**Figure 1A**). In this study, we used P-value thresholds of *P* ≤ 5 × 10^―8^to filter the GWAS significant single-nucleotide polymorphisms (SNPs). We downloaded the comprehensive GWAS summary statistics (UK Biobank eBMD and Fracture GWAS Data Release 2018) from the GEnetic Factor for Osteoporosis (GEFOS) consortium website [14]. The GEFOS consortium is an extensive international collaboration comprising numerous research groups from multiple academic institutions focusing on osteoporosis disease to accelerate gene discovery. After downloading the data, we used the statistical program R version 4.0.4 software (The R Foundation) [15]. The summary statistics of genetic association were available for 13,753,401 SNPs for eBMD. Thresholds at *P* ≤ 5 × 10^―8^ resulted in 103,155 candidate GWAS tagged SNPs and 2,955 genes, respectively.

### Functional Study with Genotype-Tissue Expression Data

In our previous study [7], we utilized the functional mapping and annotation of GWAS findings to prioritize the most likely causal SNPs and genes using information from 18 biological data repositories and tools [16]. Gene-based GWAS analysis was performed with MAGMA 1.6 [17]. EPDR1, PKDCC, and SPTBN1 genes were highly expressed in muscle tissue (**Figure 1B**).

### Single-cell Transcriptome Profiling of an Adult Human Cell in 15 Major Organs

He et al. [6] investigated the transcriptional heterogeneity and interactions of cells from an adult human’s 15 organs at the single-cell resolution level (**Figure 1C**). Using scRNA-seq technology, they profiled the transcriptomes of more than 84,000 cells of 15 organs from one individual donor. This study provides a high-resolution adult human cell atlas (AHCA), aiming at a global view of various cell populations and connections in the human body. The published AHCA data sets provide a comprehensive understanding of developmental trajectories of major cell types and identify new cell types, regulators, and key molecular events that might play important roles in maintaining the homeostasis of the human body and/or those otherwise developing into human disease. The AHCA dataset was downloaded from the Gene Expression Omnibus repository with the primary accession code GSE159929.

### Single-cell RNA Sequencing Dataset Analysis

Quality control filtering, dimensionality reduction, and clustering for cells were performed using the Seurat package [18] (version 4.1.0; https://satijalab.org/Seurat) (**Figure 1D**). We used generous quality control features with **min.cells**=2 and **min.features**=10 for each organ type of data to include as many genes and cells in this data set. Mitochondrial genes were removed with **^MT-** parameters in the “PercentageFeatures” function. Violinplot was used to plot the number of features and percentage of Mitochondrial genes. After combining 15 organ types data sets, we used the “NormalizeData” and “FindVariableFeatures” functions to normalize the data and select the number of features. The number of features was set to 2000, and ‘**vst’** was used as the selection method. We used the default parameters for the “SelectIntegrationFeatures” and “FindIntegrationAnchors” functions. Principal component analysis (PCA) and the uniform manifold approximation and projection (UMAP) [19] implemented in the “RunPCA” and “RunUMAP” functions, respectively, were used to identify deviations among cells. The number of PCs was set to 30, and **pca** was utilized in the “RunUMAP” function. The default parameter with **pca** was used in the “FindNeighbors” function. A resolution number of 0.8 was utilized in the “FindClusters” function to identify cell clusters, which were visualized using UMAP. Cell annotation was performed using the R package SingleR [20], which compares the transcriptome of every single cell to reference datasets to determine cellular identity. All analytical packages were performed in R software (version 4.0.4; https://www.r-project.org).

## Data Sharing

All data generated or analyzed during the study are included in the published paper. The AHCA dataset was downloaded from the Gene Expression Omnibus repository with the primary accession code GSE159929. The GWAS summary statistics (UK Biobank eBMD and Fracture GWAS Data Release 2018) from the GEnetic Factor for Osteoporosis (GEFOS) consortium website.

## Author Contribution

Conceptualization: Jongyun Jung.

Data curation: Jongyun Jung.

Formal analysis: Jongyun Jung.

Funding acquisition: Qing Wu.

Investigation: Jongyun Jung, Qing Wu.

Methodology: Jongyun Jung.

Project administration: Jongyun Jung, Qing Wu.

Resources: Jongyun Jung, Qing Wu.

Software: Jongyun Jung.

Supervision: Jongyun Jung, Qing Wu.

Validation: Jongyun Jung.

Visualization: Jongyun Jung.

Writing –original draft: Jongyun Jung.

Writing –review & editing: Jongyun Jung, Qing Wu.

## Acknowledgments

The authors thank the original authors of Morris et al. [3] and He et al. [6] for sharing their data in public.

## Supplemental Figure

**Figure S1.**
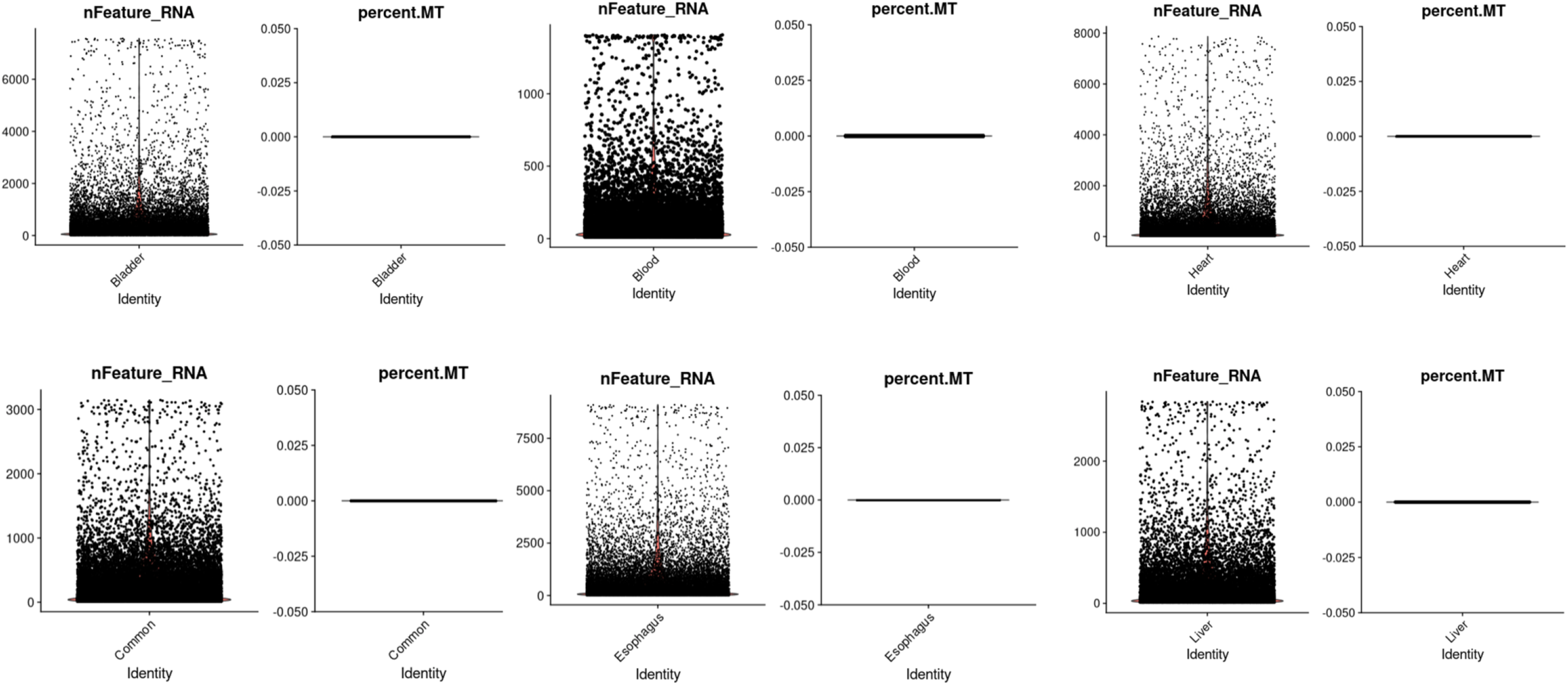
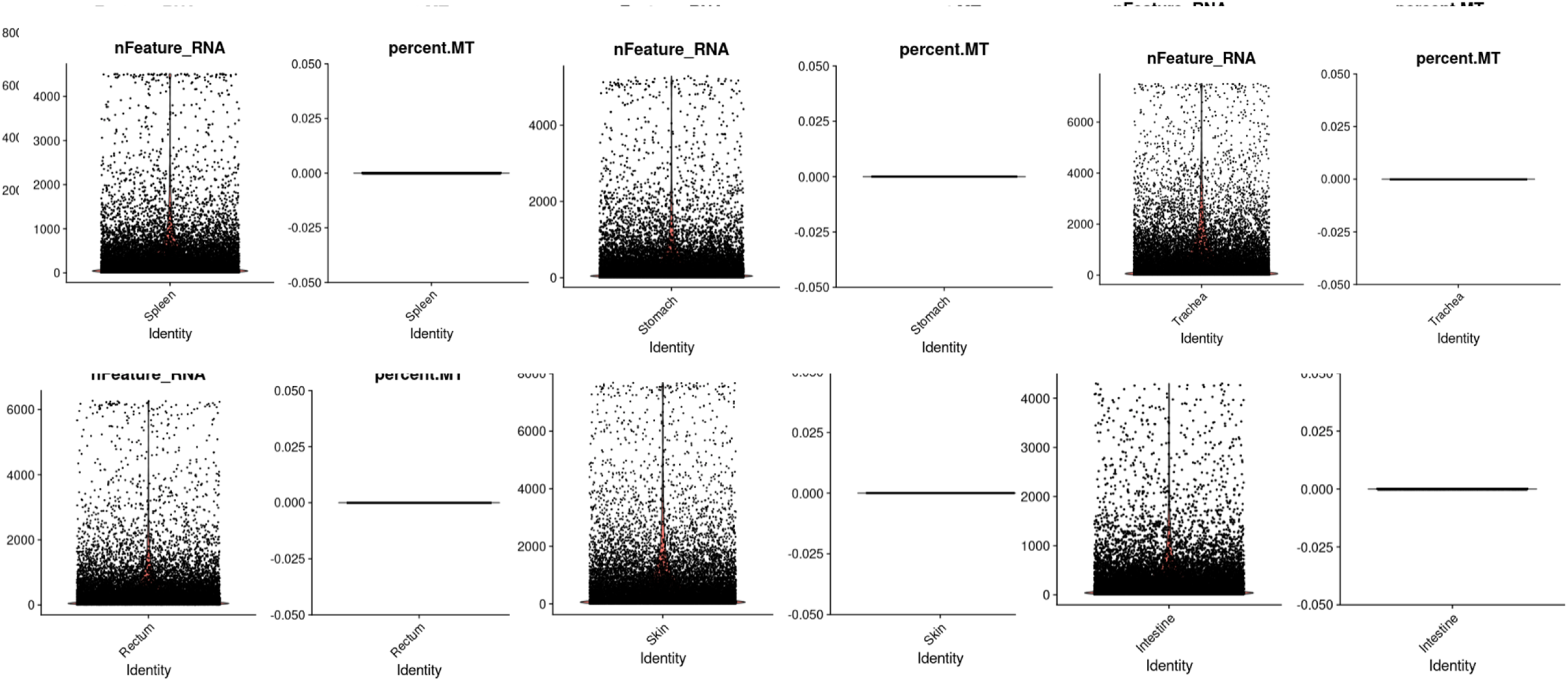
Violin plot for the quality control in each organ type.

**Figure S2.**
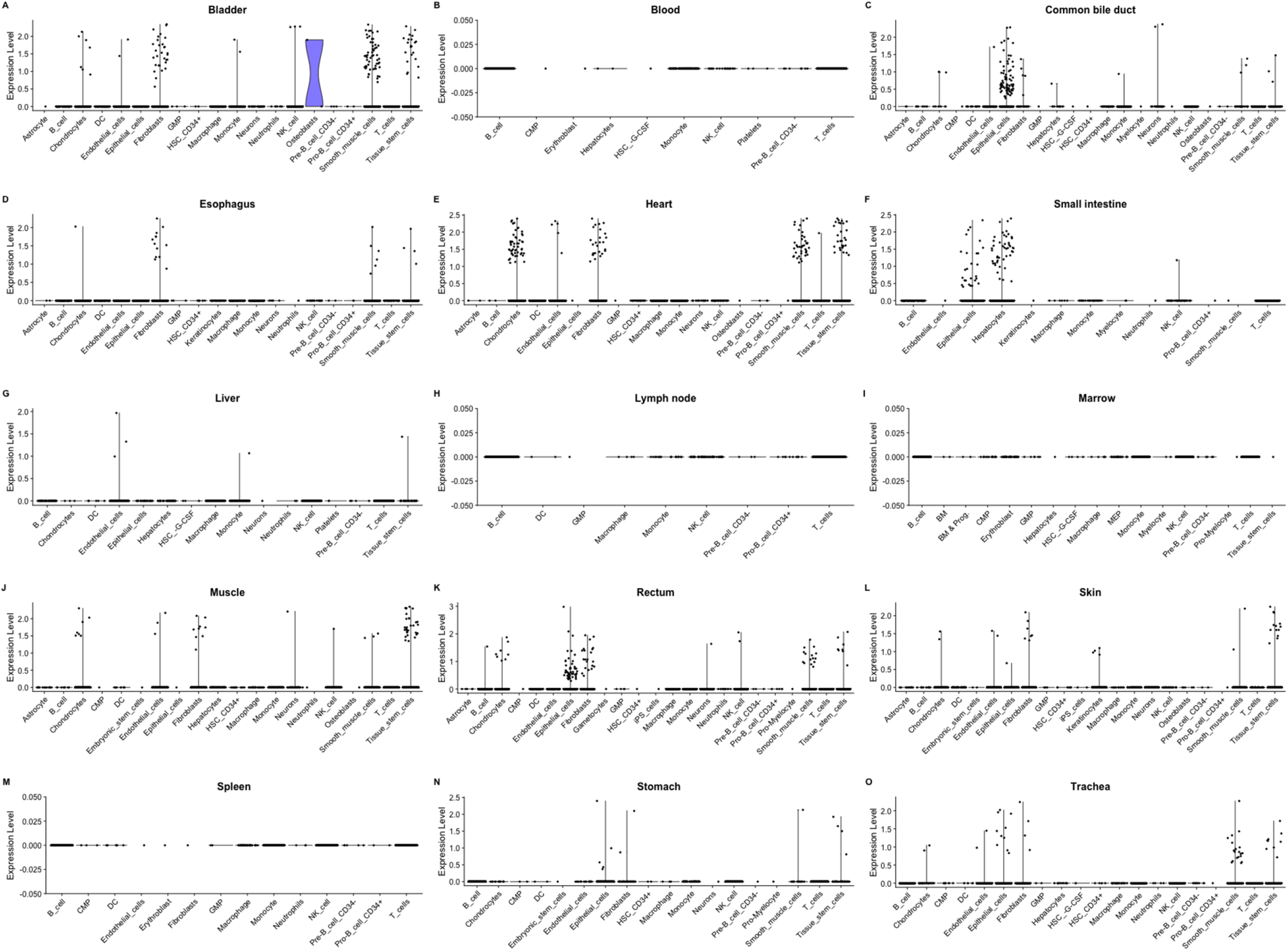
Violin plots of the normalized expression of *PKDCC* for each organ by the cell type.

**Figure S3.**
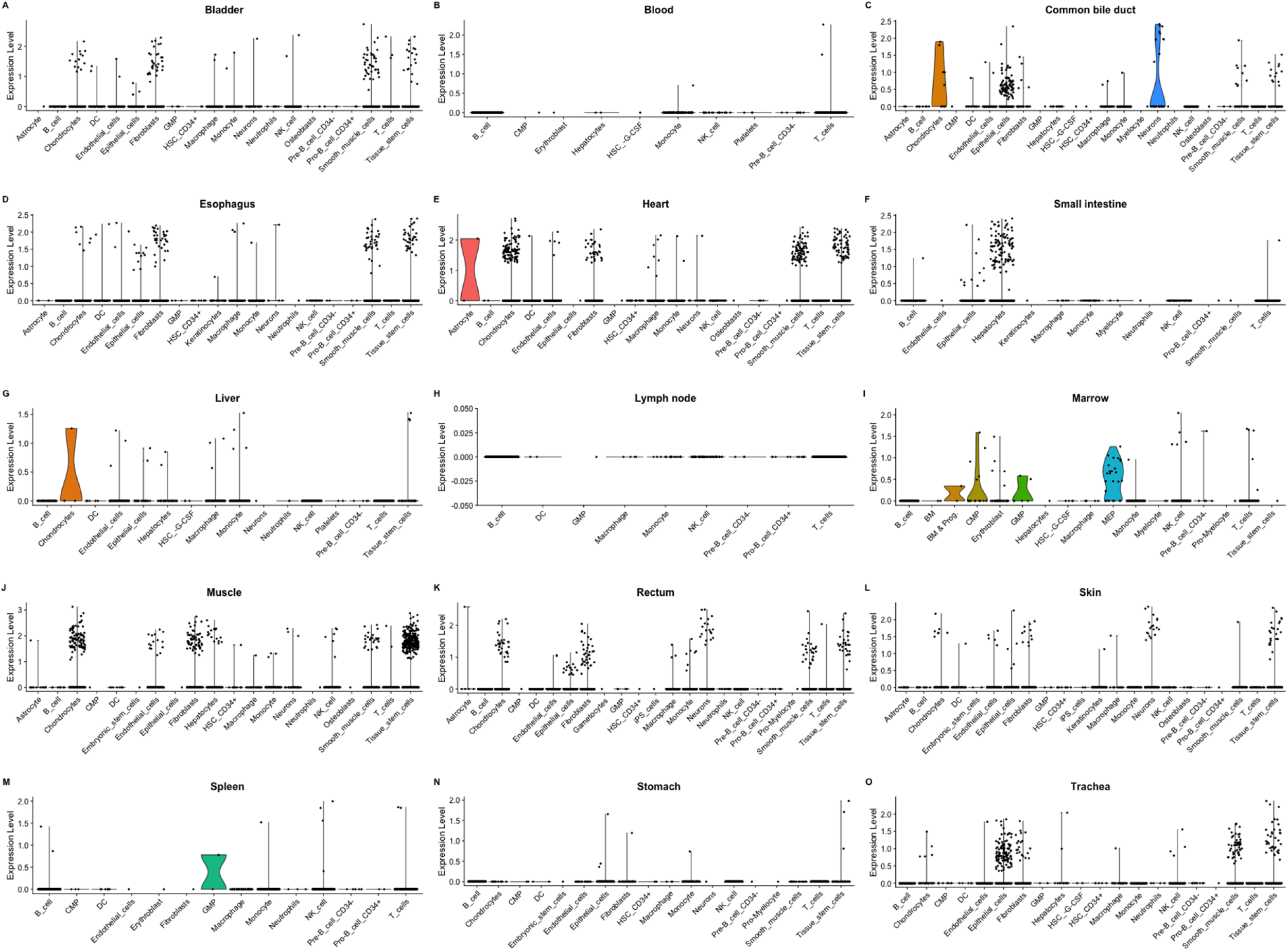
Violin plots of the normalized expression of *EPDR1* for each organ by the cell type.

